# Putative pore-forming subunits of the mechano-electrical transduction channel, Tmc1/2b, require Tmie to localize to the site of mechanotransduction in zebrafish sensory hair cells

**DOI:** 10.1101/393330

**Authors:** Itallia V. Pacentine, Teresa Nicolson

## Abstract

Mutations in transmembrane inner ear (*TMIE)* cause deafness in humans; previous studies suggest involvement in the mechano-electrical transduction (MET) complex in sensory hair cells, but TMIE’s precise role is unclear. In *tmie* zebrafish mutants, we observed that GFP-tagged Tmc1 and Tmc2b, which are putative subunits of the MET channel, fail to target to the hair bundle. In contrast, overexpression of Tmie strongly enhances the targeting of Tmc2b-GFP to stereocilia. To identify the motifs of Tmie underlying the regulation of the Tmcs, we systematically deleted or replaced peptide segments. We then assessed localization and functional rescue of each mutated/chimeric form of Tmie in *tmie* mutants. We determined that the first putative helix was dispensable and identified a novel critical region of Tmie, the extracellular region and transmembrane domain, which mediates both mechanosensitivity and Tmc2b-GFP expression in bundles. Collectively, our results suggest that Tmie’s role in sensory hair cells is to target and stabilize Tmc subunits to the site of MET.

**Author summary:** Hair cells mediate hearing and balance through the activity of a pore-forming channel in the cell membrane. The transmembrane inner ear (TMIE) protein is an essential component of the protein complex that gates this so-called mechanotransduction channel. While it is known that loss of TMIE results in deafness, the function of TMIE within the complex is unclear. Using zebrafish as a deafness model, Pacentine and Nicolson demonstrate that Tmie is required for the localization of other essential complex members, the transmembrane channel-like (Tmc) proteins, Tmc1/2b. They then evaluate twelve unique versions of Tmie, each containing mutations to different domains of Tmie. This analysis reveals that some mutations in Tmie cause dysfunctional gating of the channel as demonstrated through reduced hair cell activity, and that these same dysfunctional versions also display reduced Tmc expression at the normal site of the channel. These findings link hair cell activity with the levels of Tmc in the bundle, reinforcing the currently-debated notion that the Tmcs are the pore-forming subunits of the mechanotransduction channel. The authors conclude that Tmie, through distinct regions, is involved in both trafficking and stabilizing the Tmcs at the site of mechanotransduction.

## Introduction

The auditory and vestibular systems detect mechanical stimuli such as sound, gravity, and acceleration. These two systems share a sensory cell type called hair cells. The somas of hair cells are embedded in the epithelium and extend villi-like processes from their apex into the surrounding fluid. The shorter of these, the stereocilia, are arranged in a staircase-like pattern adjacent to a single primary cilium known as a kinocilium. Neighboring cilia are connected by protein linkages. Deflection of the kinocilium along the excitatory axis tugs the interconnected stereocilia, which move as a single unit called the hair bundle [1]. When tension is placed on the upper-most linkages known as tip links, the force is thought to open mechanosensitive channels at the distal end of the shorter stereocilia [2, 3]. These channels pass current, depolarizing the cell and permitting electrical output to the brain via the eighth cranial nerve. The conversion of a mechanical stimulus into an electrical signal is known as mechano-electrical transduction (MET) [4]. The proteins located at the site of MET and involved in gating the MET channel are collectively known as the MET complex. How the components of the MET complex, including the channel itself, are localized to and maintained at the stereocilia tips is not well understood. To characterize the molecular underpinnings of MET and the underlying cause of pathology in human patients, it is essential to examine the individual components of the transduction complex in a comprehensive fashion. Thus far, only a few proteins have been designated as members of the MET complex. The identity of the channel itself remains contentious, but currently the best candidates for the pore-forming subunits are the Transmembrane Channel-like (TMC) proteins TMC1 and TMC2. Mutations in *TMC1* cause human deafness [5], and double knock-outs of mouse *Tmc1/2* result in the loss of MET currents [6-8]. In zebrafish, overexpression of a fragment of Tmc2a generates a dominant negative effect on hair-cell mechanosensitivity [9] and Tmc2a and Tmc2b are required for MET in hair cells of the lateral line organ [10]. The TMCs localize to the tips of stereocilia, the site of MET, in mice and zebrafish [3, 6, 8, 10-12]. A point mutation in mouse *Tmc1* results in altered channel properties, suggesting direct changes to the pore [7, 13]. Likewise, in TMC2 knockout mice, channel permeation properties are altered [14]. Regardless of whether the TMCs are the pore-forming or accessory subunits of the channel, they are essential for MET.

Another key component of the complex is Protocadherin-15 (PCDH15), which comprises the lower end of the tip link [15, 16] and interacts with the TMCs [8, 9]. A fourth membrane protein, Lipoma HMGIC fusion partner-like 5 (LHFPL5, formerly called TMHS), interacts with PCDH15 and is critical for localizing PCDH15 to the site of MET [17, 18]. LHFPL5 is also required to properly localize TMC1 in mouse cochlear hair cells [8]. However, loss of LHFPL5 in cochlear hair cells does not completely abolish MET currents, and currents can be rescued by overexpression of PCDH15 [18]. This evidence suggests that LHFPL5 is not essential but rather acts as an accessory protein. Another TMC1/2 interacting partner is Calcium and integrin binding protein 2 (CIB2), which is a cytosolic protein that is localized in stereocilia and required for MET in cochlear hair cells [19].

A sixth essential member of the MET complex is the transmembrane inner ear (TMIE) protein. Loss of TMIE results in deafness in all vertebrate organisms studied [20-25]. A recent study demonstrated that TMIE is required for active MET channels in cochlear hair cells of mice [26]. These authors showed that despite normal morphology of the inner ear, hair cells lacking TMIE fail to label with aminoglycosides or FM 1-43, both of which are known to permeate the MET channel [27, 28]. TMIE was first localized to the stereocilia of hair cells [29, 30], and then to the stereocilia tips where MET occurs [25]. Zhao et al. further demonstrated that loss of TMIE ablates MET currents, that TMIE interacts with both LHFPL5 and the CD2 isoform of PCDH15, and that interfering with the TMIE-CD2 interaction alters MET. They proposed that TMIE could be a force-coupler between the tip link and channel. However, the CD2 isoform of PCDH15 is only essential in cochlear hair cells and not vestibular hair cells [31]. Zebrafish do not possess the CD2 isoform [9, 32], and yet they still require Tmie for hair-cell function [21]. These findings raised the tantalizing possibility that Tmie might have an additional role in MET that is independent from the tip links. Here, we present an alternative role for Tmie in hair cell function.

We first confirmed that mechanosensitivity is absent in a zebrafish mutant of *tmie*, *ru1000*, and demonstrated that this defect is rescued by transgenic Tmie-GFP. The localization of Tmie-GFP is maintained in the absence of other transduction components, suggesting that Tmie traffics independently to hair bundles. Unexpectedly, GFP-tagged Tmcs fail to localize to the hair bundle in *tmie* mutants, and overexpression of Tmie leads to a corresponding increase in bundle expression of Tmc2b-GFP. To determine which regions of Tmie are involved in regulating the Tmcs, we performed a domain analysis of *tmie* by expressing mutated or chimeric transgenes of *tmie* in *tmieru*^*1000*^, and made three key discoveries: (*i*) Tmie can function without its putative first transmembrane domain, (*ii*) the remaining helix (2TM) and adjacent regions are responsible for Tmie’s function in hair cells, and (*iii*) dysfunctional *tmie* constructs have reduced efficacy in localizing the Tmcs, supporting the conclusion that impaired MET is due to reduction of Tmc protein. Our evidence suggests that Tmie’s role in the MET complex is to promote localization of Tmc1/2 to the site of MET in zebrafish sensory hair cells.

## Results

### Gross morphology is normal in *tmie* ^*ru1000*^ mutant zebrafish

The literature on TMIE’s role in sensory hair cells is somewhat contradictory. Earlier studies proposed a developmental role for TMIE [20-22], while later studies evidenced a role in MET [25, 26]. To begin our analysis and attempt to clarify the issue in zebrafish, we examined live *tmie*^*ru1000*^ larvae at 5-7 dpf using confocal microscopy. The *ru1000* allele harbors a nonsense mutation leading to an N-terminal truncation, L25X [21]. We observed that mature hair cells of *tmie*^*ru1000*^ larvae were grossly normal compared to wild type siblings in both the inner ear cristae and the lateral line organ, an organ specific to fish and amphibians (Fig 1A). We noted a slight thinning of the mutant hair bundles, as revealed using a transgene Actin-GFP. Thin bundles have been observed in other zebrafish MET mutants, such as those carrying mutations in *ap1b1* and *tomt.* Both genes have been previously implicated in protein trafficking in hair cells, with *tomt* having a specific role in targeting Tmc1/2 proteins to the hair bundle [11, 33].

**Fig 1.**
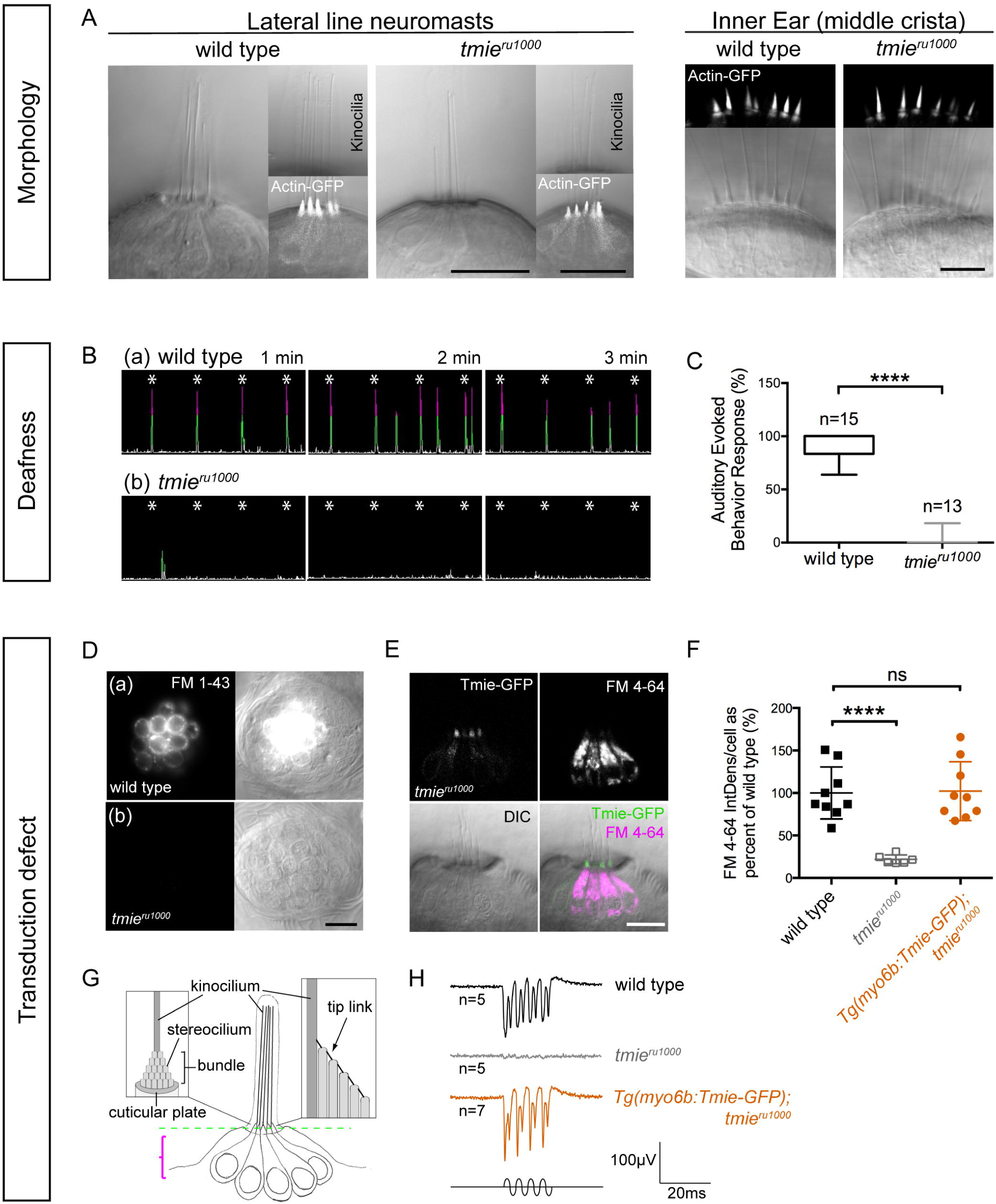
Zebrafish *tmie*^*ru1000*^ mutants: phenotype and functional rescue by Tmie-GFP. All confocal images are of live, anesthetized larvae. (A) Hair cells in the lateral-line neuromasts (7 dpf) and inner ear cristae (5 dpf) from wild type and *tmie*^*ru1000*^ larvae. A transgene (Actin-GFP) was used to visualize stereocilia bundles. (B) Sample traces from an auditory evoked behavior response (AEBR) assay, performed on 6 dpf larvae over the course of 3 minutes. Pure tone stimuli are indicated by asterisks. Peaks represent pixel changes due to larval movements (magenta indicates positive response). (C) Quantification of AEBR displayed as box-and-whiskers plot; significance determined by unpaired t-test with Welch’s correction. (D) Top-down view of neuromasts from 4 dpf larvae after brief exposure to a vital dye, FM 1-43. FM 1-43 and FM4-64 permeate open transduction channels. (E) Lateral view of a neuromast from a 4 dpf *tmie*^*ru1000*^ larva expressing transgenic Tmie-GFP, after exposure to FM 4-64. (F) Quantification of FM 4-64 fluorescence/cell in 5 dpf larvae; significance determined by one-way ANOVA. (G) A cartoon depiction of a group of lateral-line hair cells viewed laterally, with close-up views of a single cell at the bundle region. The dashed green line indicates the single plane containing the stereocilia bundles. The magenta bracket indicates the area used to make the maximum projections that were analyzed for FM fluorescence in (F). (H) Sample traces of extracellular (microphonic) recordings, evoked from the inner ear of 3 dpf larvae. A piezo actuator was used to stimulate larvae with a 200 Hz sine-wave mechanical stimulus using an 8 V driver voltage. All statistics are mean ± SD, ****p<0.0001. Scale bars: 10µm.

### Tmie-deficient zebrafish are deaf due to a defect in hair cell mechanosensitivity

Next, we used an assay for the auditory evoked behavior response (AEBR) to quantify hearing loss in *tmie*^*ru1000*^ mutants. We exposed 6 dpf larvae to a loud pure tone stimulus (157 dB, 1000 Hz, 100 ms) once every 15 seconds for three minutes and recorded their startle responses (sample traces in Fig 1B). Larvae deficient in *tmie* appear to be profoundly deaf, with little to no response as compared to wild type siblings (Fig 1B and 1C). We then determined basal (unevoked) hair cell activity of *tmie*^*ru1000*^ larvae using FM 1-43 or FM 4-64. Both are vital dyes that permeate open channels, making them useful as a proxy measure of the presence of active MET channels in hair cells [27, 28, 34]. A 30-second bath application of FM dye readily labels hair cells of the lateral line organ, which are arranged in superficial clusters called neuromasts. We briefly exposed wild type and *tmie*^*ru1000*^ larvae to FM dye and then imaged the neuromasts (Fig 1D). Consistent with previous findings [21, 22, 26], *tmie*^*ru1000*^ neuromasts have a severe reduction in FM labeling, suggesting that these hair cells have a MET defect (Fig 1F). To characterize mechanically evoked responses of hair cells, we recorded extracellular potentials, or microphonics (Fig 1H). Using a piezo actuator, we applied a 200 Hz sine wave stimulus to 3 dpf larvae while simultaneously recording voltage responses from hair cells of the inner ear. In agreement with results from our FM dye assay and with microphonic recordings previously reported [21], microphonics are absent in *tmie*^*ru1000*^ larvae (Fig 1H, gray trace).

### Transgenic *tmie-GFP* rescues the functional defect in *tmie*^*ru1000*^ mutants

To rescue mechanosensitivity in *tmie*^*ru1000*^ larvae, we generated a construct of *tmie* tagged with GFP on its C-terminus, then expressed this transgene using a hair cell-specific promoter, *myosin 6b (myo6b)*. Stably expressed Tmie-GFP rescued the FM labeling in *tmie*^*ru1000*^ hair cells (Fig 1E and 1F). Tmie-GFP also restores microphonic potentials to wild-type levels (Fig 1H, orange trace). In a stable line with a single transgene insertion, we observed that Tmie-GFP expression varies among hair cells, even within the same patch of neuroepithelium (lateral crista, S1A Fig). Immature hair cells, which can be identified by their shorter stereocilia and kinocilia (S1A Fig, bracket and arrow, respectively), consistently show a bright and diffuse pattern of labeling. This high expression level in immature bundles is characteristic of transgenes expressed using the *myo6b* promoter, which drives expression more strongly in young hair cells [17, 34]. In mature hair cells, expression patterns of Tmie-GFP are variable. At high expression levels, Tmie-GFP is enriched in the bundle in a broader pattern (S1B Fig). At reduced levels, the GFP signal is concentrated at the beveled edge of the hair bundle (S1C Fig). At very low levels, we can observe puncta along the stereocilia staircase, consistent with localization at stereocilia tips (S1D Fig). We suspect that the diffuse “bundle fill” pattern is due to overexpression, and that lower levels of Tmie-GFP recapitulate the endogenous localization at the site of MET, as previously observed in mice [25].

### Tmie-GFP is capable of trafficking without other members of the MET complex

Having confirmed that our exogenously expressed Tmie-GFP is functional, we used this transgene to probe Tmie’s role in the MET complex. First, we characterized Tmie’s interactions with other MET proteins *in vivo* by expressing transgenic Tmie-GFP in mutant *pcdh15a*, *lhfpl5a*, and *tomt* larvae (Fig 2). Because a triple knock-out of zebrafish *tmc* has not been reported, we used *tomt* mutants as a proxy for *tmc*-deficient fish based on recent studies of defective bundle localization of the Tmcs in *tomt*-deficient fish and mice [11, 35]. As in wild type bundles (Fig 2A), Tmie-GFP is detectable in the stereocilia in each of these MET mutants (Fig 2B and 2D), even if hair bundles are splayed (Fig 2B and 2C, arrowheads). This result suggests that Tmie does not depend on interactions with other MET components for entry into the hair bundle.

**Fig 2.**
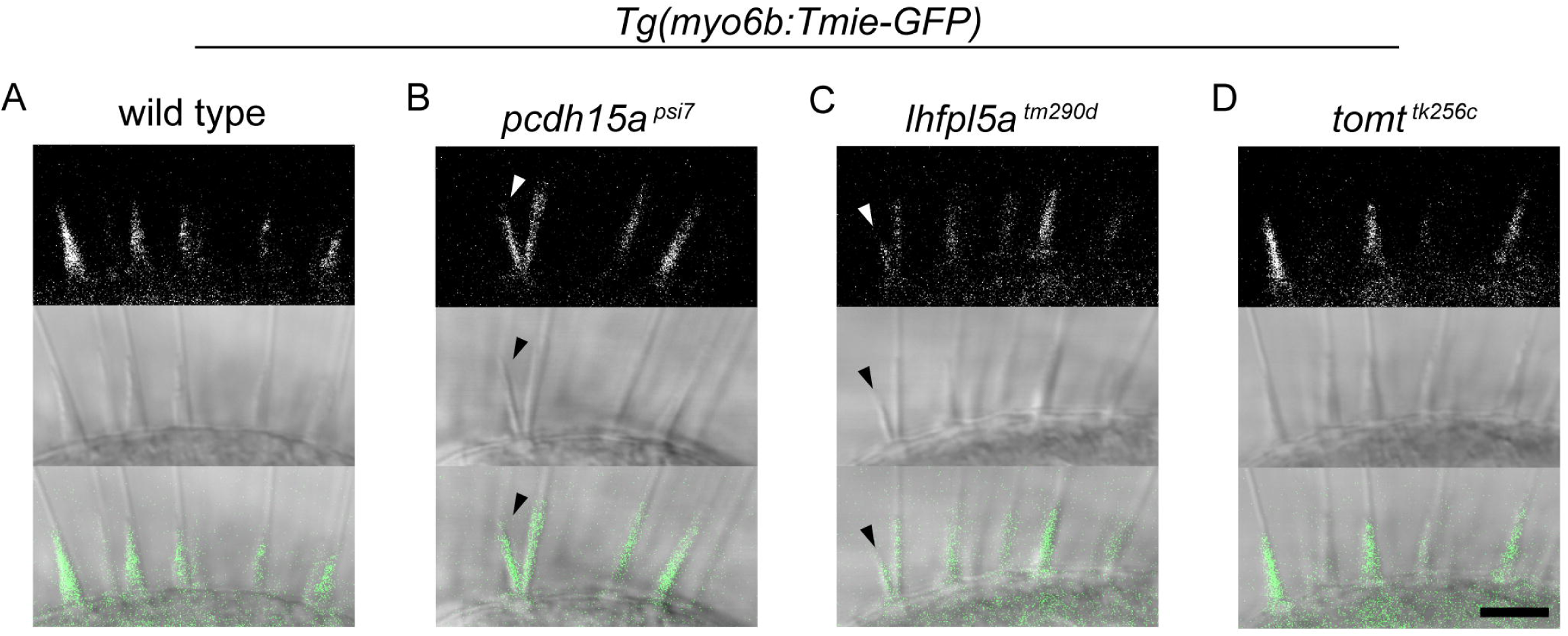
Tmie-GFP is present in the hair bundles of MET mutants. Confocal images of the bundle region in hair cells of the inner-ear lateral crista in live larvae. Larvae at 6 dpf expressing transgenic Tmie-GFP in the genetic backgrounds of wild type (A), and homozygous mutants for the tip link protein Pcdh15a (B, *pcdh15a*^*psi7*^), the accessory protein Lhfpl5a (C, *lhfpl5a*^*tm290d*^), and the Golgi-localized protein Tomt (D, *tomt*^*tk256c*^). Tomt-deficient fish lack Tmc expression in hair cell bundles [11], presumably mimicking the condition of a triple Tmc knockout. Arrowheads indicate splayed hair bundles. n=8 each genotype. Scale bar: 5µm.

### Tmc1-GFP and Tmc2b-GFP fail to localize to stereocilia without Tmie

To determine if the loss of Tmie affects the other components of the mechanotransduction complex, we expressed GFP-tagged mechanotransduction proteins (Pcdh15aCD3, Lhfpl5a, Tmc1, and Tmc2b) in *tmie*^*ru1000*^ mutants. Both Pcdh15aCD3-GFP (Fig 3A) and GFP-Lhfpl5a (Fig 3B) showed GFP fluorescence in hair bundles with a punctate distribution, similar to the pattern seen in wild type bundles. This result is consistent with the intact morphology of *tmie*^*ru1000*^ hair bundles. However, when we imaged Tmc1-GFP (Fig 3C) and Tmc2b-GFP (Fig 3E), GFP fluorescence was severely reduced in the hair bundles of *tmie*^*ru1000*^ mutants. In mature *tmie*^*ru1000*^ hair cells, we often saw a signal within the apical soma near the cuticular plate, indicative of a trafficking defect (Fig 3E, arrows; position of cuticular plate denoted in Fig 1G). We quantified Tmc expression in the hair bundle region and observed a striking and consistent reduction in *tmie* mutants (Fig 3D and 3F). Previously, we reported that localization of transgenic Tmc-GFP is unaffected in *pcdh15a* mutants [11], demonstrating that mislocalization of Tmc1/2 is not a hallmark of all MET mutants.

**Fig 3.**
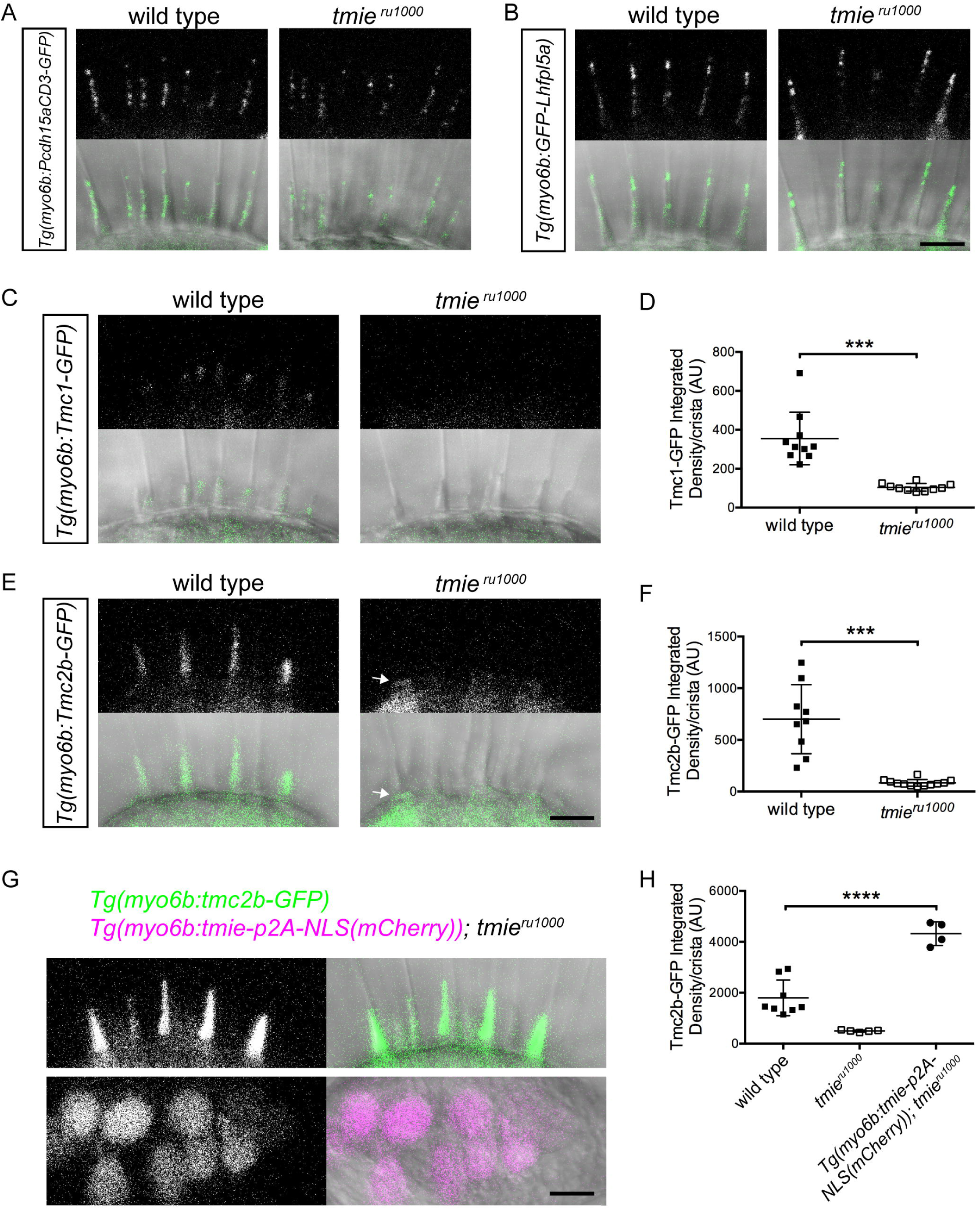
Specific loss of Tmc1 and Tmc2b in tmieru1000 larvae. Maximum projections of the hair bundle region (ROI) of hair cells in the lateral crista of the inner ear, collected from live larvae using confocal microscopy. (A-B) 6 dpf larvae expressing either transgenic Pcdh15aCD3-GFP or GFP-Lhfpl5a (n=6 each genotype). (C) 3 dpf larvae expressing Tmc1-GFP. (D) Plot of the integrated density of Tmc1-GFP fluorescence in the ROI; each data point represents one crista. Statistical significance determined by two-tailed unpaired t-test with Welch’s correction, p=0.0002. (E) 4 dpf larvae expressing Tmc2b-GFP. The arrow points to the cuticular plate/apical soma region, just below the ROI. (F) Plot of the integrated density of Tmc2b-GFP fluorescence in the ROI. Statistical significance determined by two-tailed unpaired t-test with Welch’s correction, p=0.0005. (G) 4 dpf *tmie*^*ru1000*^ larva co-expressing two transgenes, *tmc2b-GFP* and *tmie-p2A-NLS(mCherry)*. The p2A linker is a self-cleaving peptide that results in equimolar expression of Tmie and nuclear mCherry. (H) Plot of the integrated density of Tmc2b-GFP fluorescence/crista in the ROI; significance determined by one-way ANOVA. All statistics are mean ± SD. Scale bars: 5µm.

### Overexpression of Tmie increases bundle localization of Tmc2b-GFP

We hypothesized that if the loss of Tmie reduces Tmc localization in the hair bundle, then overexpression of Tmie may have the opposite effect. To test the consequence of overexpression of Tmie on Tmc localization, we created a second construct of *tmie* coupled with *p2A-NLS(mCherry)* driven by the *myo6b* promoter. The p2A linker is a self-cleaving peptide, which leads to translation of equimolar amounts of Tmie and NLS(mCherry). Hence, mCherry expression in the nucleus denotes Tmie expression in the cell (Fig 3G, lower panels). We generated a stable *tmie*^*ru1000*^ fish line carrying the *tmie-p2A-NLS(mCherry)* transgene and then crossed it to the *Tg(myo6b:tmc2b-GFP); tmie*^*ru1000*^ line. We observed that overexpression of Tmie led to a 2.5-fold increase in expression of Tmc2b-GFP in the bundles of hair cells when compared to wild type siblings that carried only the *tmc2b-GFP* transgene (Fig 3G and 3H). Combined with the finding that Tmc expression is lost in hair bundles lacking Tmie, our data suggest that Tmie positively regulates Tmc localization to the hair bundle.

### Transgenes can effectively determine protein functionality

To gain a better understanding of Tmie’s role in regulating the Tmcs, we characterized a new allele of *tmie*, *t26171*, which was isolated in a forward genetics screen for balance and hearing defects in zebrafish larvae. Sequencing revealed that *tmie*^*t26171*^ fish carry an A→G mutation in the splice acceptor of the final exon of *tmie*, which leads to use of a nearby cryptic splice acceptor (S2A Fig, *DNA, cDNA*). Use of the cryptic acceptor causes a frameshift that terminates the protein at amino acid 139 (A140X), thus removing a significant portion of the C-terminal tail (S2A Fig, *Protein*). Homozygous mutant larvae exhibit severe auditory and vestibular deficits, being insensitive to acoustic stimuli and unable to maintain balance (S2A Fig, *Balance*). FM 4-64 labeling of *tmie*^*t26171*^ mutant hair cells suggests that the effect of the mutation is similar to the *ru1000* mutation (S2B and S2D Figs). This finding implicates the C-terminal tail, a previously uncharacterized region, in Tmie’s role in MET. However, when we overexpressed a near-mimic of the predicted protein product of *tmie*^*t26171*^ (1-138-GFP) using the *myo6b* promoter, we observed full rescue of FM labeling defects in *tmie*^*ru1000*^ (S2C and S2D Figs), as well as behavioral rescue of balance and acoustic sensitivity (n=19). These results revealed that when expressed at higher levels, loss of residues 139-231 does not have a significant impact on Tmie’s ability to function.

This paradoxical finding highlighted an important advantage of the use of transgenes over traditional mutants. There are myriad reasons why a genomic mutation may lead to dysfunction, including reduced transcription or translation, protein misfolding and degradation, or mistrafficking. Exogenous expression may overcome these deficiencies by producing proteins at higher levels. Moreover, the use of transgenes enabled us to carry out a more comprehensive structure/function study of Tmie. To test a collection of deletions and chimeras of Tmie, we therefore used the *myo6b* promoter to drive exogenous expression of the constructs in hair cells of the *tmie*^*ru1000*^ mutant.

We systematically deleted or replaced regions of *tmie* to generate 12 unique *tmie* constructs (Fig 4A). Earlier studies in zebrafish and mice proposed that Tmie undergoes cleavage, resulting in a single-pass mature protein [21, 36]. To test this hypothesis, we generated the SP44-231 construct of Tmie, which replaced the N-terminus with a known signal peptide (SP) from a zebrafish Glutamate receptor protein (Gria2a). The purpose of the unrelated signal peptide was to preserve the predicted membrane topology of Tmie. We also made a similar construct that begins at amino acid 63, where the sequence of Tmie becomes highly conserved (*SP63-231*). Three of the constructs contained internal deletions (*Δ63-73; Δ97-113; Δ114-138).* In three more constructs, we replaced part of or the entire second transmembrane helix (2TM) with a dissimilar helix from the CD8 glycoprotein (*CD8; CD8-2TM; 2TM-CD8*). We included our mimic of the zebrafish *tmie*^*t26171*^ mutant, which truncates the cytoplasmic C-terminus (*1-138*). To further truncate the C-terminus, we made a construct that mimics the mouse *sr*^*J*^ mutant (*1-113*). In mice, this truncation recapitulates the full-deletion phenotype [20]. Finally, we included an alternate isoform of Tmie that uses a different final exon, changing the C-terminal sequence (*Tmie-short*). This isoform is found only in zebrafish [21] and its function has not been explored.

**Fig 4.**
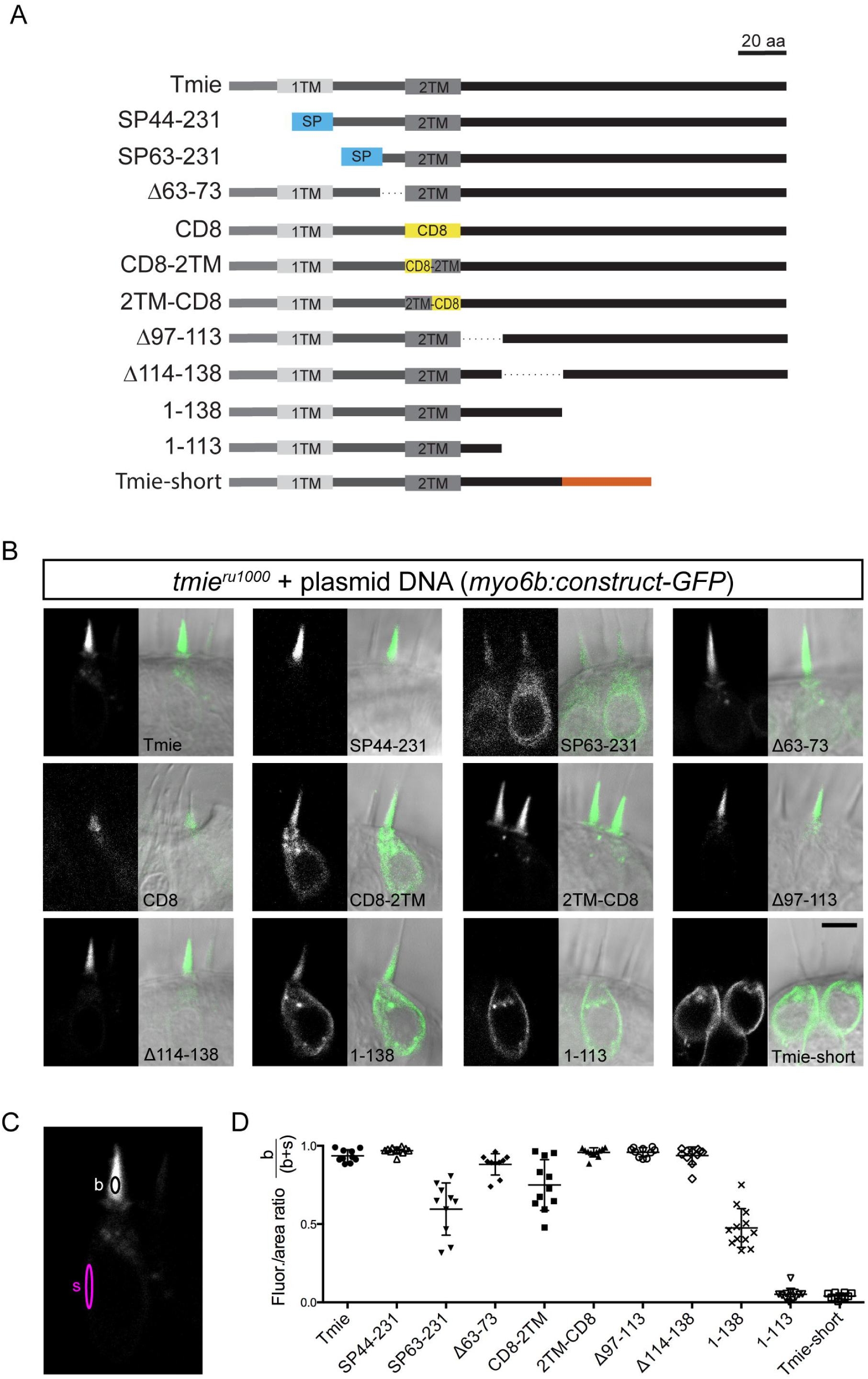
Schema for a systematic domain analysis of Tmie and subcellular localization of Tmie constructs. (A) A linear diagram of 12 unique constructs of tmie used in our experiments. Full-length Tmie is predicted to contain two hydrophobic helices or transmembrane domains (1TM and 2TM). SP44-231 and SP63-231 replace part of the N-terminus with a signal peptide (SP) from the Glutamate receptor 2a (in blue). In the CD8, CD8-2TM, and 2TM-CD8 constructs, all or part of the 2TM is replaced by the helix from the CD8 glycoprotein (in yellow). Tmie-short is a fish-specific isoform of Tmie that contains an alternate final exon (in orange). Dotted lines represent internal deletions. (B) Representative confocal images of each construct being expressed as a GFP-tagged transgene in hair cells of 4-6 dpf *tmie*^*ru1000*^ larvae. Expression is mosaic due to random genomic insertion into subsets of progenitor cells after single-cell injection. The expression of the CD8 construct is shown in a neuromast, while all others are in the inner ear middle crista. (C) The localization of each GFP fusion protein was determined by measuring the fluorescence/area in the bundle (b) and soma (s), and then calculating b/(b+s). (D) Enrichment in the hair bundle is displayed as a ratio for each construct, with 1 being completely bundle-enriched and 0 being completely soma-enriched. Scale bar in (B): 5µm.

### Subcellular localization of mutated or chimeric Tmie reveals domains required for self-localization to the bundle

We first determined the subcellular localization of each Tmie fusion protein. Plasmid DNA was co-injected into *tmie*^*ru1000*^ eggs with transposase to generate mosaic expression of the constructs in a subset of hair cells. At 4-6 days post injection, we imaged individual hair cells expressing each transgene (Fig 4B). To quantify the enrichment in the bundle versus soma, we measured the integrated density of GFP fluorescence in a small central area of mature bundles (Fig 4C, black oval) and separately in the plasma membrane or soma-enriched compartments (Fig 4C, magenta oval). Correcting for area, we then divided the bundle values by the total values (bundle/bundle + soma) and expressed this as a ratio (Fig 4D). Values closer to 1 are bundle enriched, while values closer to 0 are soma-enriched. We excluded the *CD8-GFP* construct from further analyses because it was detected only in immature bundles (Fig 4B, *CD8*).

Localization fell into three broad categories: bundle-enriched, soma-enriched, and equally distributed. Most of the fusion proteins were bundle-enriched, similar to full-length Tmie-GFP expression (Fig 4B and 4D). Three constructs were trafficked to the bundle but also expressed in the soma (*SP63-231, CD8-2TM, 1-138*). This result suggests that the deleted regions in these constructs have some role in designating Tmie as a bundle-localized protein. Also of note, the full replacement of the 2TM helix (*CD8*) was unable to maintain stable expression in mature bundles. Half-TM replacements (*CD8-2TM, 2TM-CD8*) revealed that loss of the first half of the helix affects trafficking, whereas alteration of the second half had no effect. Only two constructs were soma-enriched (*Tmie-short* and *1-113*), suggesting an inability to traffic to the bundle. These two transgenes were thus excluded from further analyses.

### FM labeling identifies functional regions in the second transmembrane domain and adjacent residues of Tmie

To identify regions of Tmie involved in mechanosensitivity of hair cells, we measured the functionality of the nine *tmie* constructs that showed hair bundle expression. As in Fig 1F, we generated stable lines of each transgenic construct and quantified fluorescence in neuromasts after exposure to FM 4-64 (Fig 5).

**Fig 5.**
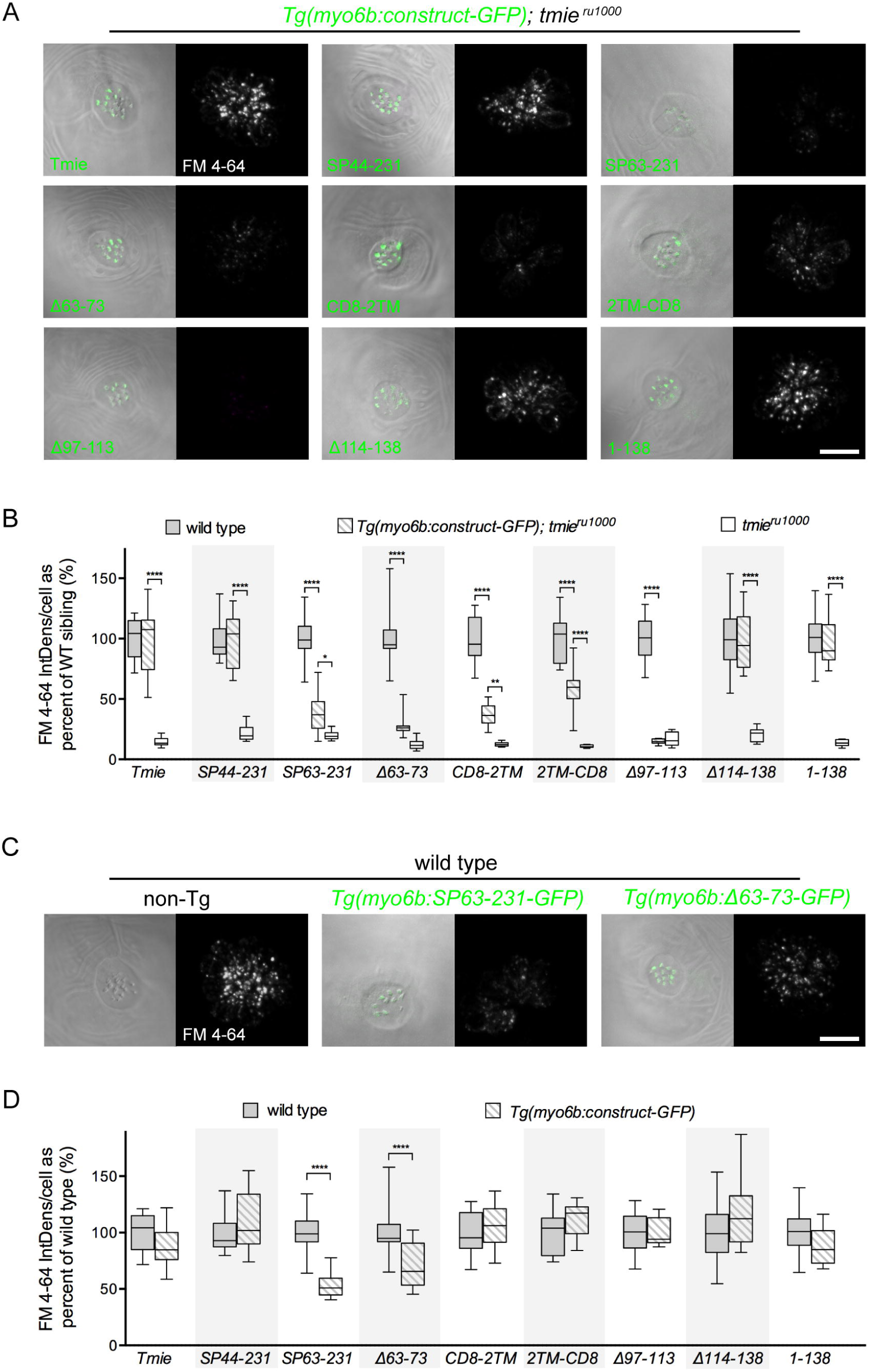
The second transmembrane and adjacent residues of Tmie are required for rescue of FM labeling. All images are a top-down view of a representative neuromast from 6 dpf larvae collected using confocal microscopy. The left image is a single plane through the stereocilia (green dashed line in Fig 1G) with DIC + GFP fluorescence. The right image is a maximum projection of the soma region (magenta bracket in Fig 1G) showing FM 4-64 fluorescence. (A) Representative images of neuromasts in *tmie*^*ru1000*^ larvae, each stably expressing an individual tmie construct. FM fluorescence was normalized to wild type non-transgenic larvae generated with the Tmie-GFP line. (B) Box-and-whiskers plot of the integrated density of FM fluorescence/cell in transgenic *tmie*^*ru1000*^ compared to non-transgenic wild type and mutant siblings for each construct. (C) Representative images of neuromasts in wild type larvae with or without transgene. FM fluorescence was normalized to wild type non-transgenic larvae of the Tmie-GFP line. (D) Box-and-whiskers plot of the integrated density of FM fluorescence/cell in wild type neuromasts with and without transgene. Significance determined within each clutch by one-way ANOVA, n≥9, **p<0.01, ***p<0.001, ****p<0.0001. Scale bars in (A) and (C) are 10µm.

Of nine constructs examined, four showed wild type levels of FM fluorescence in *tmie*^*ru1000*^ neuromasts (*Tmie, SP44-231, Δ114-138*, and *1-138;* Fig 5A and 5B). Two constructs (*Δ97-113* and *Δ63-73*) did not rescue above mutant levels of FM 4-64, although *Δ*63-73 showed a non-significant increase in FM fluorescence. While residues 63-73 have not been characterized, the *Δ*97-113 result is consistent with the findings of previous publications in humans and mice, showing that mutations in this region impair hearing and hair cell function [23, 25]. Three constructs were capable of partial rescue (*SP63-231, CD8-2TM,* and *2TM-CD8*). Each one of the five dysfunctional constructs altered part of a contiguous region of Tmie: the 2TM and surrounding domains. These results highlight this region of Tmie as vital for function. To determine whether any of the constructs also have a dominant effect on hair-cell function, we compared FM label in wild type larvae with or without the individual transgenic *tmie* construct (Fig 5D). SP63-231 and *Δ*63-73, which had impaired rescue in *tmie*^*ru1000*^, showed reduced FM label in transgenic wild type cells (Fig 5C and 5D). Interestingly, these two dominant negative constructs alter the extracellular region of Tmie.

### Recordings of mechanically evoked responses confirm that the second transmembrane domain and adjacent regions are required for hair cell function

Bath applied FM dye demonstrates the presence of permeable MET channels, but does not reveal any changes in mechanically evoked responses in hair cells. Therefore, we also recorded microphonics of mutant larvae expressing individual transgenes. For our recordings, we inserted a recording pipette into the inner ear cavity of 3 dpf larvae and pressed a glass probe against the head (Fig 6A). Using a piezo actuator to drive the probe, we delivered a step stimulus at increasing driver voltages while recording traces in current clamp (Fig 6B). For each transgenic *tmie* line, we measured the amplitude of the response at the onset of stimulus (Fig 6C-I). We limited our analysis to the lines expressing constructs that failed to fully rescue FM labeling (Fig 6E-I). As positive controls, we used the full-length *tmie* line (Fig 65 C) and also included *SP44-231* (Fig 6D), encoding the cleavage product mimic. Both controls fully rescued the responses in *tmie*^*ru1000*^ larvae. Consistent with a reduction in labeling with FM dye, we found that the microphonic responses were strongly or severely reduced in larvae expressing the *SP63-231, Δ63-73, CD8-2TM*, *2TM-CD8* and *Δ97-113* constructs in the *tmie*^*ru1000*^ background (Fig 6E-I). We also saw the same dominant negative effect in wild type larvae expressing transgenic *SP63-231* or *Δ63-73.*

**Fig 6.**
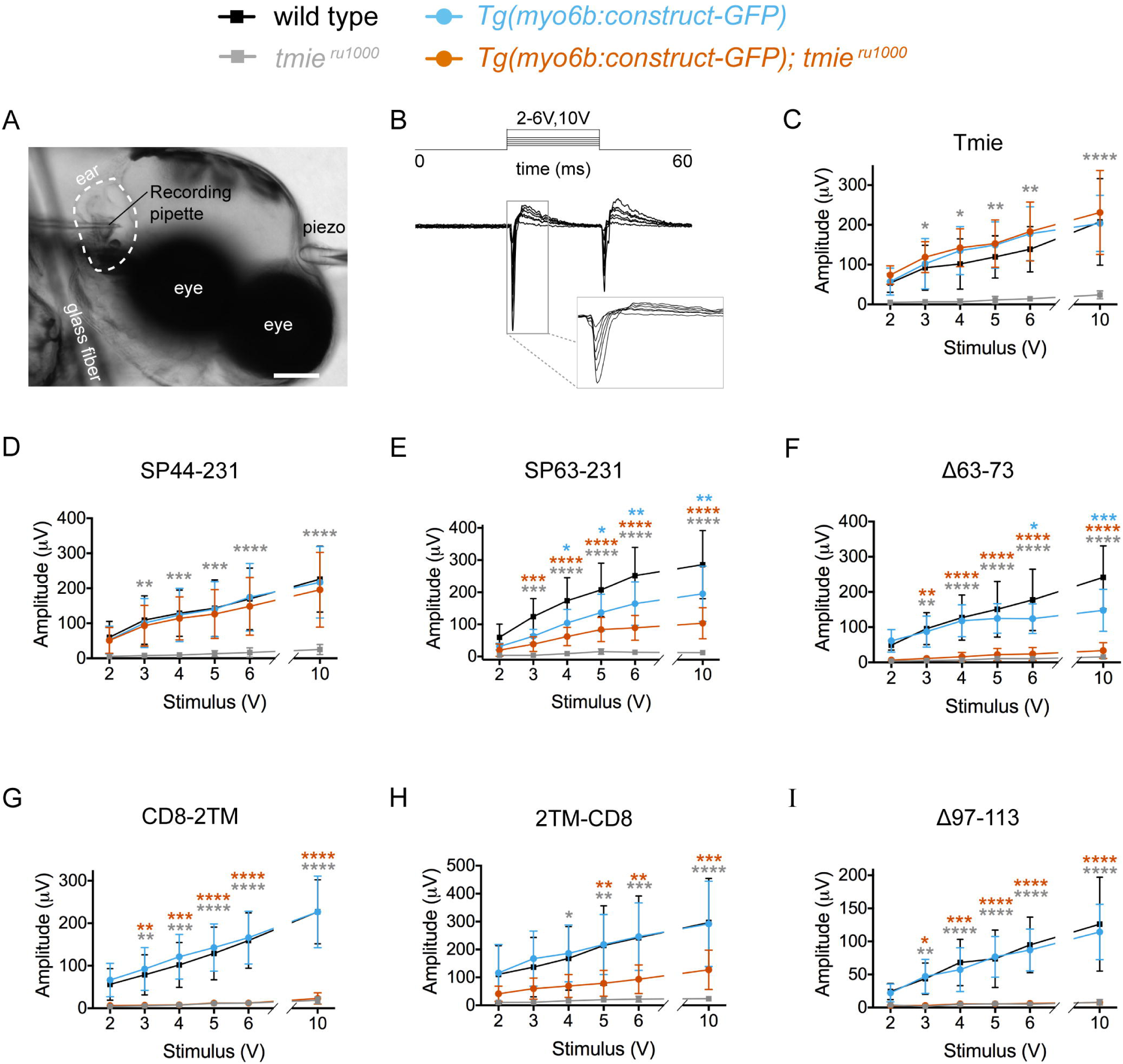
The second transmembrane and adjacent regions of Tmie are required for inner ear microphonics. (A) A DIC image of a 3 dpf larva anesthetized and pinned (glass fiber) for inner ear recordings. Shown are a probe attached to a piezo actuator (piezo) pressed against the head and a recording pipette pierced into the inner ear. (B) Traces from a wild type larva. A step stimulus for 20ms was applied; 200 traces were averaged for each of the six piezo driver voltages: 2V, 3V, 4V, 5V, 6V, and 10V. Gray box: magnification of the onset of response in individual traces. (C-I) Same protocol as in (B). Mean amplitude of the response peak ± SD as a function of the stimulus intensity of the driver voltage. Statistical significance determined by two-way ANOVA comparing all groups to wild type non-transgenic siblings, n≥5, *p<0.05, **p<0.01, ***p<0.001, ****p<0.0001. Scale bar: 100µm.

### Regions of Tmie that mediate hair-cell mechanosensitivity are also required for localizing Tmc2b-GFP

After identifying functional regions of Tmie, we asked whether these regions are involved in regulating Tmc localization. Therefore we quantified hair bundle expression of transgenic Tmc2b-GFP in hair cells of *tmie*^*ru1000*^ mutant larvae stably co-expressing individual transgenic *tmie* constructs (Fig 7B-H). As in Fig 3 G, we tagged our *tmie* constructs with p2A-NLS(mCherry) so that Tmc2b-GFP expression in the hair bundles could be imaged separately. We examined SP44-231 and the five *tmie* constructs that yielded impaired mechanosensitivity.

**Fig 7.**
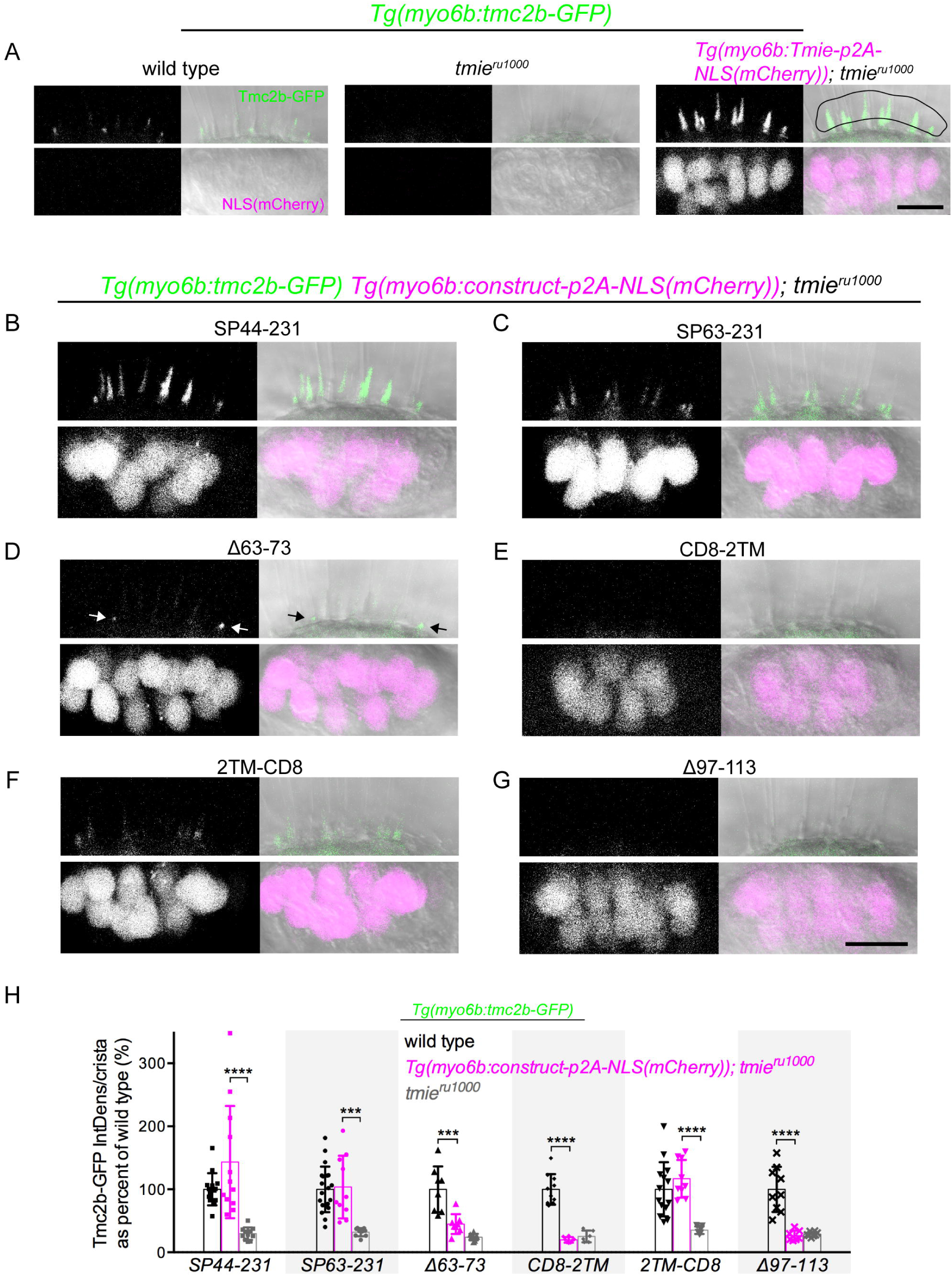
Effect of transgenic Tmie constructs on Tmc2b-GFP bundle localization. Confocal images are maximum projections of representative inner-ear lateral cristae collected from 4 dpf larvae. Upper panels show the bundle region, with all larvae stably expressing transgenic Tmc2b-GFP (green). Lower panels show the soma region, with some larvae expressing transgenic Tmie constructs tagged with p2A-NLS(mCherry). Nuclear mCherry (magenta) is a marker for equimolar translation of the indicated Tmie construct. (A) Sibling wild type, *tmie*^*ru1000*^, and *tmie*^*ru1000*^ expressing full-length Tmie. For the quantification in (G), Tmc2b-GFP fluorescence was measured within the ROI (right panel, black line). (B-G) *tmie*^*ru1000*^ larvae expressing individual Tmie constructs tagged with p2A-NLS(mCherry), as labeled. The arrows in (D) point to Tmc2b-GFP in immature hair bundles. (H) Plot of the integrated density of Tmc2b-GFP fluorescence in the ROI, comparing *tmie*^*ru1000*^ larvae expressing a tmie construct (magenta) to wild type (black) and *tmie*^*ru1000*^ (gray) siblings not expressing tmie construct. Significance for SP44-231 and SP63-231 was determined by the Kruskal-Wallis test, for all other tmie constructs by one-way ANOVA, n≥6, ***p<0.001, ****p<0.0001. Scale bars: 10µm.

Three constructs showed full rescue of Tmc2b-GFP levels in the bundle. The SP44-231 cleavage mimic produced highly variable levels, some in the wild type range, others increasing Tmc2b-GFP expression above wild type (Figs 7B and 7H), as seen in overexpression of full-length Tmie (Figs 3H and 7A, right panel). We suspect that the exogenous Gria2a signal peptide leads to variable processing of Tmie and thus contributes to this variability in Tmc2b-GFP fluorescence. *tmie*^*ru1000*^ larvae expressing the SP63-231 construct gave rise to values of Tmc2b-GFP fluorescence within the wild type range (Fig 7C and 7H). When we recorded microphonics in these larvae, we found that co-overexpression of Tmc2b-GFP and SP63-231 resulted in better functional rescue of *tmie*^*ru1000*^ (S3A Fig) than when SP63-231 was expressed alone (Fig 6E). We also determined that the microphonic potentials correlated with the levels of Tmc2b-GFP in the bundles (S3B Fig). Likewise, the 2TM-CD8 construct also generated values of Tmc2b-GFP fluorescence in the wild type range (Fig 7F and 7H). These larvae rescued microphonic potentials to wild type levels (S3C Fig), unlike when 2TM-CD8 was expressed alone (Fig 6E). Functional rescue again correlated with Tmc2b-GFP bundle levels (S3D Fig). These results indicate that functional rescue in the SP63-231 and 2TM-CD8 lines is Tmc dose-dependent.

Of the three constructs with little to no functional rescue, CD8-2TM (Fig 7E and 7H) and *Δ*97-113 (Fig 7G and 7H) had severely reduced levels of Tmc2b-GFP in hair bundles. In *tmie*^*ru1000*^ expressing *Δ*63-73, there was severely reduced but still faintly detectable Tmc2b-GFP signal though, as with the functional rescue, this difference was not statistically significant (Fig 7D and 7H). The bulk of this signal was observed in immature bundles (Fig 7D, arrows, and S4 Fig), but there was some detectable Tmc2b-GFP signal in mature bundles (S4 Fig). Overall, these results suggest that the level of functional rescue by the *tmie* constructs is correlated to the amount of Tmc2b present in the hair bundle.

## Discussion

*TMIE* was first identified as a deafness gene in mice and humans [20, 23]. The predicted gene product is a relatively small membrane protein containing a highly conserved amino acid sequence near the second hydrophobic helix. Previous studies established that TMIE is required for MET in hair cells [21, 25, 26] and is an integral member of the complex (Zhao et al., 2014). How TMIE contributes to the function of the MET complex was not clear. Our comprehensive structure-function analysis of Tmie revealed that the functional capacity of various *tmie* mutant constructs is determined by their efficacy in localizing Tmc2b-GFP to the bundle, as summarized in Fig 8A and modeled in Fig 8B. These findings unveil a hitherto unexpected role for Tmie in promoting the localization of the putative channel subunits Tmc1 and Tmc2b to the site of MET. These findings broaden our understanding of the assembly of the MET complex and point to a pivotal role of Tmie in this process.

**Fig 8.**
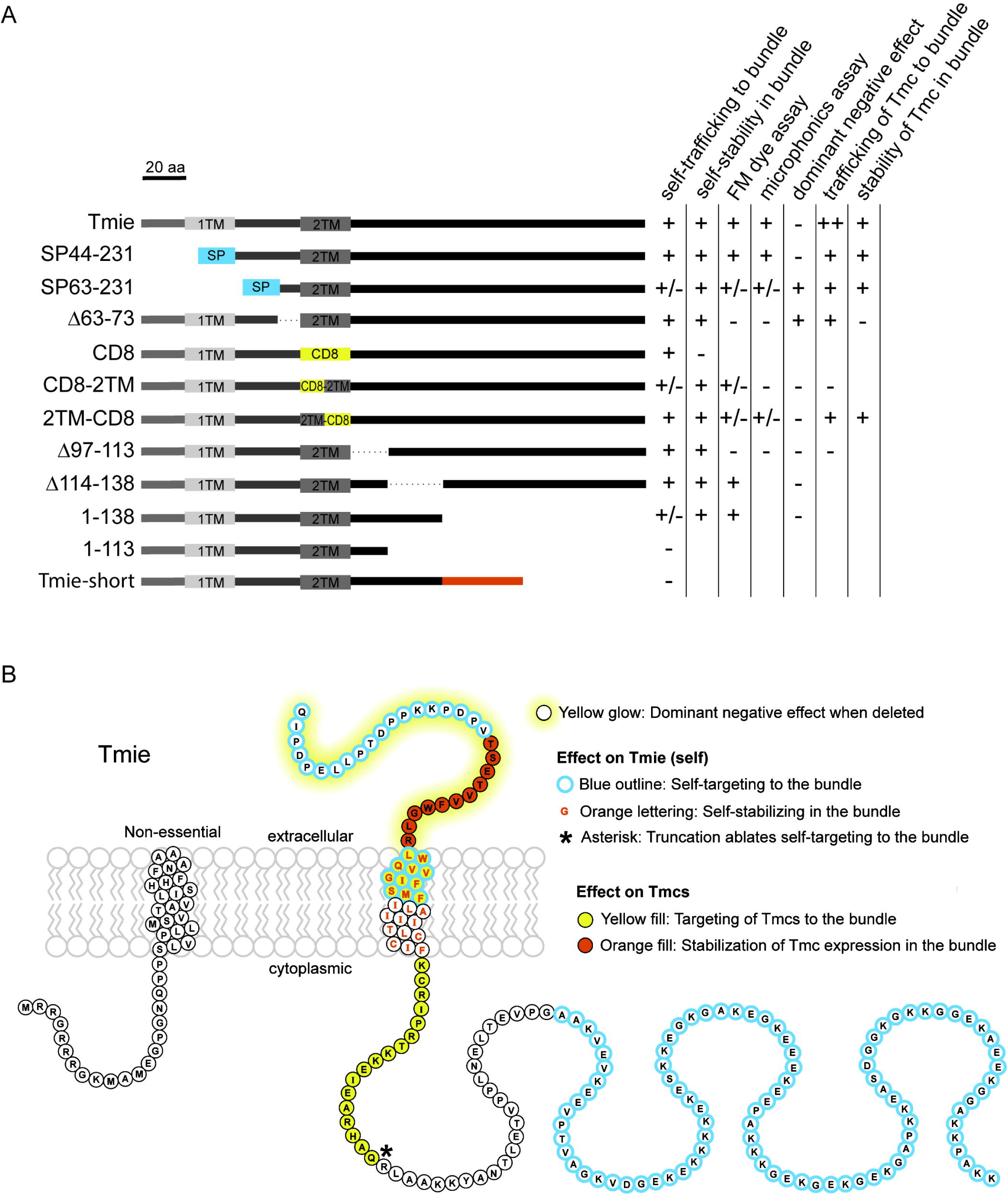
Summary of experimental results for tmie constructs and model of discrete functional domains of Tmie. (A) Symbols as follows: enhanced above wild type (++), comparable to wild type (+), partially reduced (+/-), and severely reduced or absent (-). For *dominant negative effect*, the effect is present (+) or absent (-). Blank spaces are not determined. Refer to Fig 4A for details on constructs. (B) A model of the protein sequence of zebrafish Tmie. Amino acids 1-43 are separated due to suspected cleavage as a signal peptide. Although shown as a TM domain, it is unclear whether the first hydrophobic region forms a helix. Note that the extracellular region (yellow glow) was never deleted in its entirety; SP63-231 deleted QIPDPELLPTDPPKKPDPV, and *Δ*63-73 deleted TSETVVFWGLR. Also note that the TM domain with orange lettering only had an effect on the stability of Tmie when the entire helix was substituted.

A previous study of the *ru1000* mutant suggested that Tmie’s role in zebrafish hair cells was developmental, with mutant lateral line hair cells showing stunted kinocilia and the absence of tip links [21]. In our hands we did not observe any gross morphological defects, and the localization pattern and the levels of Pcdh15a and Lhfpl5a were unaffected in *tmie*^*ru1000*^ larvae. This observation is consistent with intact hair bundle morphology; stereocilia that are splayed or disorganized are a dominant feature of hair cells missing their tip links, as seen in *pcdh15a* or *lhfpl5a* mutants [17, 32]. In TMIE-deficient mice, hair cell morphology is grossly normal up to P7 [25, 26]. In agreement with a previous study in mice [25], our results indicate that *tmie*^*ru1000*^ mutants are profoundly deaf due to ablation of MET in hair cells. We fully rescued this deficit in zebrafish *ru1000* mutants by exogenous expression of a GFP-tagged transgene of *tmie*. Exogenous expression gave rise to variable levels of Tmie-GFP in hair bundles, with lower levels revealing a punctate pattern expected for a member of the MET complex, and higher expression levels leading to expression throughout the stereocilia. Excess Tmie-GFP did not appear to cause adverse effects in hair cells, which is consistent with a previous study in the *circler* mouse mutant [37].

### Tmie can localize to hair bundles independently

To determine the interdependence of trafficking of Tmie and the other MET components, we examined the localization of Tmie-GFP in mutants of essential MET genes: the tip-link protein *pcdh15a*; the accessory protein *lhfpl5a*; and in *tomt* mutants (Fig 3). In both zebrafish and mice, the secretory pathway protein Tomt is required for proper localization of the Tmc1/2 [11, 35]; the *tomt* mutant likely simulates the condition of a triple knockout of all three zebrafish *tmc* genes (*tmc1/2a/2b*). Despite the absence of Pcdh15a, Lhfpl5a, and Tmcs, we found that Tmie-GFP still traffics to the bundles of hair cells. This finding suggests that Tmie can localize independently of the other proteins of the MET apparatus. This autonomy is an unusual feature for membrane components of the MET complex. For example, PCDH15 largely requires LHFPL5 for trafficking to the stereocilia [17, 18, 38], and depends on Cadherin 23 to maintain its localization at the site of MET [17, 39]. LHFPL5 also requires PCDH15 to maintain localization at the stereocilia tips [18, 38]. Thus, Tmie appears to be the exception to the rule of co-dependent transport to the hair bundle.

Our results reveal that Tmie has distinct regions associated with self-localization and function (Fig 8). Three constructs showed impaired targeting of Tmie to the bundle, namely SP63-231, CD8-2TM, and the 1-138 construct, the last of which truncates the C-terminus. Further manipulation to the C-terminus, either by removing more amino acids (*1-113*) or by using an alternative final exon (*Tmie-short*), results in targeting of the protein to the plasma membrane instead of the bundle. However, removing just a smaller internal segment has no effect on bundle localization (Fig 4A and 4C, *Δ114-138*). We suspect that the abundance of charged residues in the C-terminus of Tmie (S2A Fig, *Protein*), as well as the regions altered in SP63-231 and CD8-2TM, contribute to recognition by bundle trafficking machinery. Mislocalization, however, did not necessarily correlate with functional rescue. Despite partial mislocalization to the plasma membrane, the 1-138 construct showed full functional rescue. Conversely, despite normal localization to the bundle, Δ97-113 did not rescue function at all. These results demonstrate that Tmie’s functional role is separate from its ability to target to the bundle.

### Tmie promotes the levels of Tmc1/2 in the hair bundle

The regulatory role of Tmie with respect to the Tmcs is strongly supported by the strikingly different effects of loss of Tmie versus overexpression of Tmie. When Tmie is absent, so are the Tmcs; when Tmie is overexpressed, the level of Tmc2b in the bundle is boosted as well (Fig 3C-H). These results disagree with a previous finding in mice showing that Myc-TMC2 is present in hair bundles of TMIE-deficient cochlear hair cells [25]. This discrepancy may be due to the use a cytomegalovirus promoter to drive high levels of expression of Myc-TMC2 in an *in vitro* explant of cochlear tissue. Localization of TMC1 in *Tmie-/-* mice, which is the predominant TMC protein in cochlear hair cells, was not reported. In addition, localization of the TMCs in vestibular hair cells was not characterized in *Tmie-/-* mice. Thus, further investigation is warranted to determine if the relationship between Tmie and the Tmcs uncovered by our experiments is a conserved feature or is potentially dependent on the type of hair cell, as MET components may vary among different cell types.

One important question is whether Tmie and the Tmcs can physically interact to form a complex that is transported to the hair bundle. A direct interaction of the mouse TMC1/2 and TMIE proteins was not detected in a heterologous system [25], however, our *in vivo* analysis suggests the possibility of an indirect interaction. The deletions and chimeric forms of Tmie in the present study highlight important motifs or regions of Tmie that are critical for localization of the Tmcs to the hair bundle.

### The first hydrophobic helix of Tmie is dispensable

The membrane topology of Tmie has not been biochemically determined, however, online *Phobius* software predicts an N-terminal signal peptide in mouse and human TMIE and a transmembrane helix in zebrafish Tmie [40]. Interestingly, the orthologues in C. elegans or Drosophila do not contain this first hydrophobic region of Tmie. Upon removal of this region, we observed that SP44-231 behaved like full-length Tmie, with a comparable pattern of localization and full functional rescue of *tmie*-deficient fish. In addition, SP44-231 rescues Tmc2b-GFP bundle expression to wild type levels or higher. To our knowledge, these results are the first *in vivo* evidence that Tmie can function without the putative first transmembrane domain. Our study supports the notion that Tmie undergoes cleavage, resulting in a single-pass membrane protein that functions in the MET complex (Fig 8B).

### The 2TM domain and adjacent regions of Tmie are functionally significant

The key functional domains of Tmie are located within and proximal to the remaining transmembrane domain. We found that replacement of the entire transmembrane domain with an exogenous membrane helix from the CD8 glycoprotein resulted in a protein that trafficked to the bundles of immature hair cells but was not expressed in mature bundles. This finding demonstrates that this domain is vital for stable localization of Tmie in mature hair cells. Half-chimeras of this domain revealed that the mislocalization effect is exclusive to the first half of the helix, but that both halves are functionally significant (the first half more so than the second). These results suggest that the transmembrane domain is critical for both Tmie’s localization and function in the MET complex.

Removal of the cytoplasmic amino acids 97-113, directly after the 2TM, leads to a normal localization pattern but complete loss of function. This region contains arginine residues that have previously been implicated in human deafness [23, 41-43]. Mimics of these mutations in mouse cochlear hair cells lead to altered MET currents, which has been attributed to a reduction in binding to PCDH15-CD2 [25]. Interestingly, one of the mouse mutations, R93W, resulted in loss of TMIE localization at the site of MET. In contrast to these findings, when we remove this entire intracellular region from zebrafish Tmie, it is still capable of localization in hair bundles. This result may reflect species differences in recognition sequences for trafficking machinery.

The SP63-231 and Δ63-73 constructs both lack different segments of the extracellular region of Tmie. These were the only two constructs with dominant negative effects, suggesting that each construct successfully integrates into the MET complex and interferes or competes with endogenous Tmie. Both constructs only minimally rescue mechanosensitivity in *tmie*^*ru1000*^ mutants and are thus predicted to weaken the efficiency of the MET complex. Other constructs such as the transmembrane chimeras also yield partial rescue but do not appear to affect the function of endogenous Tmie in wild-type hair cells. These data suggest that the full 2TM domain is required to produce the dominant negative effect on endogenous Tmie. Combined with the finding that replacement of the 2TM with an unrelated helix causes instability of Tmie in mature hair cells, we suggest that the 2TM is essential in integrating Tmie into the MET complex.

### Impaired functionality corresponds to decreased Tmc expression

When co-expressed with Tmc2b-GFP, our Tmie constructs reveal a strong link between function and Tmc bundle expression (Figs 7 and S3). In larvae expressing CD8-2TM and Δ97-113, both of which display little or no functional rescue, there is no detectable Tmc2b-GFP in the hair bundle (Fig 7E, 7G and 7H). In addition to defects in targeting Tmcs to the hair bundle, our data also suggest a role for Tmie in maintaining the levels of Tmc2b in stereocilia. The most dramatic effect on maintenance of Tmc signal in the bundle was seen in *tmie*^*ru1000*^ larvae expressing the Δ63-73 construct. In these larvae, Tmc2b-GFP successfully traffics to the bundle in immature hair cells (Fig 7D, arrows) but does not maintain strong expression in mature cells (S4 Fig). Based on this data, we conclude that the first half of the transmembrane domain and the intracellular residues 97-113 are involved in trafficking the Tmcs to the site of MET, while the extracellular residues 63-73 stabilize Tmc expression in the MET complex (Fig 8B).

Surprisingly, SP63-231 and 2TM-CD8 rescue Tmc2b-GFP to wild type levels (Fig 7C, 7F and 7H), even though functional rescue of *tmie*^*ru1000*^ by GFP-tagged versions was reduced in both FM labeling experiments (Fig 5) and microphonic recordings of the inner ear (Fig 6). This result hints at an additional role for Tmie in MET that is independent of Tmc trafficking. However, the low level of functional rescue in *tmie*^*ru1000*^ mutants by these two constructs was only observed in the background of endogenous levels of the Tmcs. When we co-expressed Tmc2b-GFP with either SP63-231 or 2TM-CD8, then the functional rescue of *tmie*^*ru1000*^ mechanosensitivity improved in a Tmc-dose-dependent manner (S3 Fig). Since co-expression of Tmc2b-GFP can overcome the functional deficit in SP63-231 and 2TM-CD8, we propose that residues 44-62 and the second half of the 2TM are important but not absolutely essential to regulating Tmc bundle expression. This finding reinforces the significance of our findings with the constructs Δ63-73, CD8-2TM, and Δ97-113, which still fail to rescue Tmc2b-GFP levels even when Tmc2b-GFP is co-expressed.

Through a systematic *in vivo* analysis of *tmie* via transgenic expression, we identified new functional domains of Tmie. We demonstrated a strong link between Tmie’s function and Tmc1/2 expression in the bundle. Evidence continues to mount that the Tmcs are subunits of the MET channel, and our results implicate Tmie in promoting and maintaining the localization of Tmc subunits at the site of MET. The precise mechanism underlying Tmie’s regulation of the Tmcs awaits further investigation.

## Methods

### Zebrafish husbandry

Zebrafish (*Danio rerio*, txid7955) were maintained at 28°C and bred according to standard conditions. All animal research was in compliance with guidelines from the Institutional Animal Care and Use Committee at Oregon Health and Science University. In this study, the following zebrafish mutant lines were used: *tmie*^*ru1000*^ [21], *tmie*^*t26171*^, *pcdh15a*^*psi7*^ [9], *lhfpl5a*^*tm290d*^ [44], *tmc2b*^*sa8817*^ [11]. All zebrafish lines in this study were maintained in a Tübingen or Top long fin wild type background. We examined larvae at 4-7 days post-fertilization (dpf), of undifferentiated sex. For experiments involving single transgenes, non-transgenic *tmie*^*ru1000*^ heterozygotes were crossed to transgenic fish in the homozygous or heterozygous *tmie*^*ru1000*^ background. Mutants were genotyped by PCR and subsequent digestion or DNA sequencing. Primers are listed in Table 1.

**Table 1.**
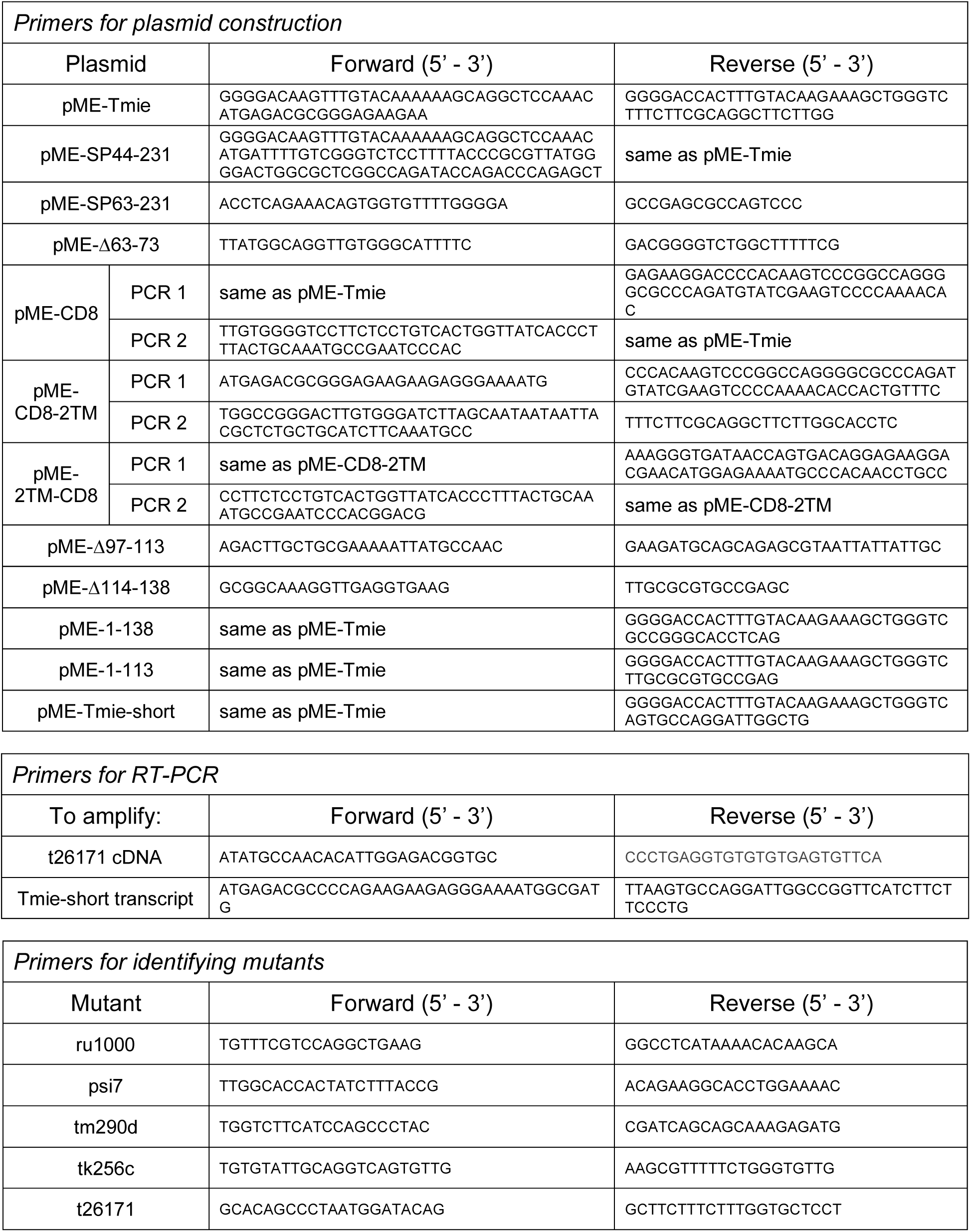
List of primers used in this study.

### Gene accession numbers for mutants and transgenes

*tmie* (accession no. F1QA80), *tmc1* (accession no. F1QFU0), *tmc2b* (accession no. F1QZE9), *tomt* (accession no. A0A193KX02), *pcdh15a* (accession no. Q5ICW6), *lhfpl5a* (accession no. F1Q837), *actba* (accession no. Q7ZVI7).

### Transgenic lines and plasmid construction

The following previously published transgenic lines were used: *Tg(-6myo6b:β-actin-GFP-pA)* [34], *Tg(-6myo6b:pcdh15aCD3-mEGFP-pA)* [17], and *Tg(-6myo6b:GFP-lhfpl5a-pA), Tg(-6myo6b:Tmc1-mEGFP-pA), Tg(-6myo6b:Tmc2b-mEGFP-pA)* [11].

To generate the *tmie* expression vectors, we used the Tol2/Gateway system [45]. The pDestination vector contained either a *cmlc2:GFP* heart marker or α-*ACry:mCherry* eye marker for sorting. pDESTtol2pACrymCherry was a gift from Joachim Berger and Peter Currie (Addgene plasmid # 64023, [46]).

The 5’ entry vector contained the promoter for the *myosin 6b* gene, which drives expression only in hair cells. All *tmie* transgenic constructs were subcloned into the middle entry vector using PCR or bridging PCR and confirmed by sequencing. The primers for each vector are listed in Table 1. For GFP-tagging, we used a 3’ entry vector with a flexible linker (GHGTGSTGSGSS) followed by *mEGFP*. For *NLS(mCherry)* experiments, a p2A self-cleaving peptide (GSGATNFSLLKQAGDVEENPGP) was interposed between the *tmie* construct and the *NLS(mCherry)*. This causes translation of a fusion protein that is subsequently cleaved into the two final proteins. The 2TM helix replacements from residues 21-43 result in the following chimeric helices: CD8 (YIWAPLAGTCGVLLLSLVITLYC), CD8-2TM (YIWAPLAGTCGILAIIITLCCIF), and 2TM-CD8 (LWQVVGIFSMFVLLLSLVITLYC).

Multisite Gateway LR reactions [47, 48] were performed to generate the following constructs: *pDest(-6myo6b:tmie-GFP-pA)*, *pDest(-6myo6b:tmie-short-GFP-pA), pDest(-6myo6b:SP44-231-GFP-pA), pDest(-6myo6b:SP63-231-GFP-pA), pDest(-6myo6b:Δ63-73-GFP-pA), pDest(-6myo6b:CD8-GFP-pA), pDest(-6myo6b:CD8-2TM-GFP-pA), pDest(-6myo6b:2TM-CD8-GFP-pA), pDest(-6myo6b:Δ97-113-GFP-pA), pDest(-6myo6b:Δ114-138-GFP-pA), pDest(-6myo6b:1-113-GFP-pA), pDest(-6myo6b:1-138-GFP-pA), pDest(-6myo6b:tmie-p2A-NLS(mCherry)-pA), pDest(-6myo6b:SP63-231-p2A-NLS(mCherry)-pA), pDest(-6myo6b:Δ63-73-p2A-NLS(mCherry)-pA), pDest(-6myo6b:CD8-2TM-p2A-NLS(mCherry)-pA), pDest(-6myo6b:Δ97-113-p2A-NLS(mCherry)-pA).*

To generate transgenic fish, plasmid DNA and *tol2* transposase mRNA were co-injected into single-cell fertilized eggs, as previously described (Kwan et al., 2007). For each construct, 200+ eggs from an incross of *tmie*^*ru1000*^ heterozygotes were injected. To obtain stable transgenic lines, >24 larvae with strong marker expression were raised as potential founders. For each GFP-tagged transgene, at least two founder lines were generated and examined for visible bundle expression. For each *tmie* construct, we isolated a line containing single transgene insertions, with the exception of the *CD8-2TM* construct in which we identified a single founder with high transmission of the transgene (>10%) and used these offspring and their siblings for FM and microphonics experiments. For *NLS(mCherry)* experiments, injected fish were raised to adulthood and genotyped to identify *tmie*^*ru1000*^ heterozygotes and homozygotes. We identified founders for each construct and then crossed these founders to *tmie*^*ru1000*^ heterozygotes carrying *Tg(myo6b:tmc2b-GFP)*. This generated offspring that expressed both transgenes in the *tmie*^*ru1000*^ mutant background, and we used these larvae for experiments. In *SP44-231*, *SP63-231*, and *CD8-2TM*, stable transgenic lines were generated from the founder before experiments were carried out.

### Microscopy

Live larvae were anesthetized with E3 plus 0.03% 3-amino benzoic acid ethylester (MESAB; Western Chemical) and mounted in 1.5% low-melting-point agarose (Sigma-Aldrich cas. # 39346-81-1), with the exception of the morphology images from Fig 1A and Fig 6A in which larvae were pinned with glass rods and imaged in E3 or extracellular solution containing MESAB. The image in Fig 6A was captured at room temperature using a Hamamatsu digital camera (C11440, ORCA-flash2.8), MetaMorph Advanced NX software, and an upright Leica DMLFS microscope. We used differential interference contrast (DIC) with a Leica HC PL Fluotar 10x/0.3 lens. For all imaging except Fig 6A, images were captured at room temperature using an Axiocam MrM camera, Zeiss Zen software, and an upright Zeiss LSM700 laser-scanning confocal microscope. We used DIC with one of two water-immersion lenses: Plan Apochromat 40x/1.0 DIC, or Acroplan 63x/0.95 W. Laser power and gain were unique for each fluorophore to prevent photobleaching. We averaged 2 or 4x for each image, consistent within each experiment. The Tmc1-GFP and Tmc2b-GFP transgenes are very dim, and high laser power (4%) and gain (1100) were necessary. At these settings, autofluorescence from other wavelengths can falsely enhance the emission peak at 488. To filter out this autofluorescence, we simultaneously collected light on a second channel with an emission peak at 640 nm.

### Auditory Evoked Behavioral Response (AEBR)

Experiments were conducted as previously described [49]. Wild type and mutant larvae were sorted by FM 1-43 labeling. Briefly, 6 dpf larvae were placed in six central wells of a 96-well microplate mounted on an audio speaker. Pure tones were played every 15 s for 3 min (twelve 100 ms stimuli at 1 kHz, sound pressure level 157 dB, denoted by asterisks in Fig 1B). Responses were recorded in the dark inside a Zebrabox monitoring system (ViewPoint Life Sciences). Peaks represent pixel changes from larval movement. A response was considered positive if it occurred within two seconds after the stimulus and surpassed threshold to be considered evoked, not spontaneous (Fig 1B, green indicates movement detected, magenta indicates threshold surpassed). For each larva, we used the best response rate out of three trials. Response was quantified by dividing the number of positive responses by total stimuli (12) and converting to a percent. If the larvae moved within two seconds before a stimulus, that stimulus was dropped from the trial data set (i.e. the number of total stimuli would become 11). Each data point on the graph in Fig 1C is the percent response of an individual larva. We used a two-tailed unpaired t-test with Welch’s correction to determine significance, ****p<0.0001.

### FM 1-43 and FM 4-64 labeling

Larvae were briefly exposed to E3 containing either 3µM N-(3-Triethylammoniumpropyl)-4-(4-(Dibutylamino)styryl)Pyridinium Dibromide (FM 1-43, Life Technologies) or 3µM of the red-shifted *N[scap]*-(3-triethylammoniumpropyl)-4-(6-(4-(diethylamino)phenyl)hexatrienyl)pyridinium dibromide (FM4-64; Invitrogen). After exposure for 25-30 seconds, larvae were washed 3x in E3. Laser power was adjusted for each experiment to avoid saturation of pixels but was consistent within a clutch. FM levels were quantified in ImageJ [50] as described previously [9]. In brief, maximum projections of each neuromast were generated using seven optical sections, beginning at the cuticular plate and moving down through the soma (magenta bracket, Fig 1G). We then measured the integrated density of the channel with an emission peak at 640 nm for FM 4-64, and at 488 nm for FM 1-43. This integrated density value was divided by the number of cells, thus converting each neuromast into a single plot point of integrated density per cell (IntDens/cell). Statistical analyses were always performed between direct siblings. For Fig 5, individual values were divided by the mean of the sibling wild type neuromasts in order to display the data as a percent of wild type, making it easier to compare across groups. Statistical significance was determined within an individual clutch using one-way ANOVA.

### Microphonics

Larvae at 3 dpf were anesthetized in extracellular solution (140mM NaCl, 2mM KCl, 2mM CaCl_2_, 1mM MgCl_2_, and 10mM 4-(2-hydroxyethyl)-1-piperazineethanesulfonic acid (HEPES); pH 7.4) containing 0.02% 3-amino benzoic acid ethylester (MESAB; Western Chemical). Two glass fibers straddled the yolk to pin the larvae against a perpendicular cross-fiber. Recording pipettes were pulled from borosilicate glass with filament, O.D.: 1.5 mm, O.D.: 0.86 mm, 10 cm length (Sutter, item # BF150-86-10, fire polished). Using the Sutter Puller (model P-97), we pulled the pipettes into a long shank with a resistance of 10-20MΩ. We then used a Sutter Beveler with impedance meter (model BV-10C) to bevel the edges of the recording pipettes to a resistance of 3-6 MΩ. We pulled a second pipette to a long shank and fire polished to a closed bulb, and then attached this rod to a piezo actuator (shielded with tin foil). The rod was then pressed to the front of the head behind the lower eye, level with the otoliths in the ear of interest, to hold the head in place while the recording pipette was advanced until it pierced the inner ear cover. Although it has been demonstrated that size of response is unchanged by entry point [51], we maintained a consistent entry point dorsal to the anterior crista and lateral to the posterior crista (see Fig 6A). After the recording pipette was situated, the piezo pipette was then moved back to a position in light contact with the head. We drove the piezo with a High Power Amplifier (piezosystem jena, System ENT/ENV, serial # E18605), and recorded responses in current clamp mode with a patch-clamp amplifier (HEKA, EPC 10 usb double, serial # 550089). Each stimulus was of 20 ms duration, with 20 ms pre- and post-stimulus periods. We used either a sine wave or a voltage step and recorded at 20 kHz, collecting 200 traces per experiment. In Fig 1H, we used a 200 Hz sine wave at 10V, based on reports that 200 Hz elicited the strongest response [52]. In Fig 6, we used multiple step stimuli at varying voltages (2V, 3V, 4V, 5V, 6V, and 10V). The piezo signal was low-pass filtered at 500Hz using the Low-Pass Bessel Filter 8 Pole (Warner Instruments). Microphonic potential responses were amplified 1000x and filtered between 0.1-3000 Hz by the Brownlee Precision Instrumentation Amplifier (Model 440). We used Igor Pro for analysis. We averaged each set of 200 traces to generate one trace response per fish, then measured baseline-to-peak amplitude. These amplitudes were used to generate the graphs in Fig 6. Statistical significance was determined by 2-way ANOVA comparing all groups to wild type non-transgenic siblings.

### Quantification of Tg(myo6b:Tmc1-GFP) and Tg(myo6b:Tmc2b-GFP) in the ROI

Using ImageJ, maximum projections of each crista were generated for analysis (5 sections per stack for Tmc1-GFP in Fig 3D and Tmc2b-GFP in Fig 3F, and 13 sections per stack for Tmc2b-GFP in Figs 3H and 7). Quantification of Tmc-GFP bundle fluorescence in Fig 3 was achieved by outlining each bundle to encompass the entire region of interest (ROI) in a single hand-drawn area (Fig 7A, right panel, black outline). In the ROI, we quantified the integrated density of the channel with an emission peak at 480 nm. This was repeated in the region above the bundles containing only inner ear fluid and the kinocilia in order to subtract background fluorescence. Each middle crista generated one data point on the graphs in Figs 3 and 7. In some cases, we saw single cells that appeared to have a GFP-fill, probably due to clipping of the GFP tag. We excluded these cells from analyses, since they falsely increased the signal. Due to the 3D nature of the mound-shaped cristae, it was difficult to completely exclude the apical soma region, leading the signals of *tmie*^*ru1000*^ to average above zero. We used the Kruskal-Wallis test for the SP44-231, SP63-231, and 2TM-CD8 constructs; all others are one-way ANOVA.

### cDNA generation by Reverse Transcription Polymerase Chain Reaction (RT-PCR)

We sorted 30 wild type and 30 *t26171* larvae by behavior (tap sensitivity and balance defect at 5 dpf) and extracted RNA using the RNeasy mini kit (Qiagen). Larvae were homogenized using a 1ml syringe. To generate the cDNA for the short isoform of Tmie (Tmie-short) and the t26171 allele, we performed RT-PCR on these RNA samples using the RNA to cDNA EcoDry Premix (Clontech, Cat # 639549). Primers are listed in Table 1. Both transcripts were verified by DNA sequencing.

## Acknowledgements

The authors thank Cecilia Toro, Lucille Moore, and Andre Dagostin for help with execution and analysis of the microphonics experiments, as well as Larry Trussell, Josef Trapani, and Anthony Ricci for advice. We thank Jim Hudspeth for the *tmie*^*ru1000*^ fish line. We also thank Eliot Smith for feedback on the manuscript, and Leah Snyder and Lisa Hiyashi for laboratory support.

## Supporting information

**S1 Fig. Tmie-GFP shows variable expression in stereocilia.** Representative images of the lateral crista in a wild type larva at 6 dpf, generated using confocal microscopy. (A) The hair bundle region of hair cells expressing transgenic tmie-GFP driven by the myo6b promoter. The arrow and bracket show, respectively, the short kinocilium and stereocilia bundle of an immature hair cell. (B) A single hair bundle with “bundle fill” expression pattern produced by overexpression of Tmie-GFP. (C) A single bundle with Tmie-GFP concentrated along the beveled edge of the stereocilial staircase. (D) A single bundle with punctate expression of Tmie-GFP suggestive of localization at the site of MET. Scale bar in (A): 5µm, in (D): 2µm.

**S2 Fig. Differential effects on function with a genomic mutation and a transgene mimic.** (A) Data for a novel mutant allele of tmie, *t26171*. DNA: Chromatographs of the DNA sequence of tmie in wild type (above) and *tmie*^*t26171*^ (below) showing the genomic region where the mutation occurs. An arginine is mutated to guanine in the splice acceptor (black box, above) of the final exon of tmie, exon 4. The dashed black box below indicates the mutated original splice acceptor site. Use of a cryptic splice acceptor (black box, below) 8 nucleotides downstream causes a frameshift and an early stop codon (*). cDNA: Chromatograph of the DNA sequence from RT-PCR of *tmie*^*t26171*^ larvae bridging exons 3 and 4. Protein: The predicted protein products, shown here as a two-pass transmembrane protein. The wild type protein has many charged residues (positive in light gray, negative in dark gray) that are lost in *tmie*^*t26171*^. Balance: Photos of wild type and *tmie*^*t26171*^ larvae, taken with a hand-held Canon camera. Arrow points to a larva that is upside-down, displaying a classic vestibular phenotype. (B) Top-down view of a representative neuromast after exposure to FM 4-64, imaged using confocal microscopy. The first panel is a single plane through the soma region while the second panel is a maximum projection of 7 panels through the soma region, beginning at the cuticular plate (as denoted by magenta bracket in Fig 1G). (C) Same as (B) except that the first panel shows the bundle region so that 1-138-GFP can be visualized in bundles (as depicted by dashed green line, Fig 1G). The transgene is driven by the myo6b promoter. (D) Plot of the integrated density of FM fluorescence per cell as a percent of wild type siblings. Displayed wild type and *tmie*^*ru1000*^ data are from siblings of *Tg(1-138-GFP); tmie*^*ru1000*^, while *tmie*^*t26171*^ data are from a separate experiment. Statistical significance determined by one-way ANOVA, ****p<0.0001. Scale bar: 10µm.

**S3 Fig. Functional rescue of tmieru1000 by constructs SP63-231 and 2TM-CD8 is Tmc dose-dependent.** (A) Mean amplitude of the response peak ± SD as a function of the stimulus intensity of the driver voltage, as described in Fig 6B. (B) XY plot of the amplitude of microphonic response vs the integrated density of Tmc2b-GFP fluorescence in the ROI. A 10V step stimulus was used to evoke microphonic potentials. The line is a linear regression, R^2^= 0.5216. (C) Same as (A) for the 2TM-CD8 construct. (D) Same as (B) for the 2TM-CD8 construct, R^2^= 0.7726. Measurements are from 4 dpf larvae.

**S4 Fig. In Δ63-73, Tmc2b-GFP traffics to bundles but does not maintain high expression in mature cells.** Confocal images of single hair bundles from cells expressing transgenic Tmc2b-GFP driven by the myo6b promoter. Brackets show the stereocilia bundle, which is shorter in immature hair cells (white brackets) and longer in mature ones (black brackets). Larvae are 4 dpf. Scale bar: 2µm.

